# Utility of Shrimp and *Der p 10* specific IgE for Shrimp Allergy Diagnosis in Non-House Dust Mite Sensitized Patients

**DOI:** 10.1101/199620

**Authors:** Karen Thursday S. Tuano, Sara Anvari, Imelda Celine Hanson, Joud Hajjar, Filiz Seeborg, Lenora M. Noroski, Danielle Guffey, Grace Kang, Jordan Scott Orange, Carla M. Davis

## Abstract

**Background:** There are no set specific IgE (sIgE) to predict shrimp allergy as cross-reactivity with other arthropods play a role in shrimp sensitization.

**Objective:** This study identifies the allergens associated with shrimp allergy in house dust mite (HDM) and non-HDM sensitized patients.

**Methods:** Patients with shrimp sensitization (positive skin prick test [SPT] and/or sIgE) with/without history of clinical reaction were recruited. Allergy was confirmed by oral food challenge (OFC) except for patients with history of anaphylaxis. Shrimp allergic (SA) and shrimp tolerant (ST) patients were further classified based on HDM sensitivity. The sIgE to shrimp, shrimp and HDM components were performed. Fisher’s exact test, Wilcoxon sum rank test and receiver operating characteristics analyses were done.

**Results:** Of 79 patients recruited, 12 SA (7 positive OFC and 5 with history of anaphylaxis), 18 ST and 10 non-shrimp sensitized controls (NC) were enrolled. In non-HDM sensitized patients, sIgE to shrimp (10.5 kU_A_/L, p=0.012) and *Der p 10* (4.09 kU_A_/L, p=0.035) were higher in SA patients. Shrimp sIgE ≥3.55 kU_A_/L had 100% sensitivity and 85.71% specificity (ROC=0.94[0.81, 1.0]). *Der p 10* sIgE ≥3.98 kU_A_/L had sensitivity of 80% and specificity of 100% (ROC=0.86[0.57, 1.0]). *rPen a* 1 ≥1.1 kU_A_/L had sensitivity of 80% and specificity of 85.7% (ROC=0.80[0.47,1.0]).

**Conclusions:** In non-HDM sensitized patients, shrimp sIgE ≥3.55 kUA/L and Der p 10 sIgE≥3.98 kUA/L give 100% sensitivity and specificity, respectively, to diagnose shrimp allergy. HDM sensitivity can influence sIgE levels to shrimp and shrimp/HDM components due to cross-reactivity.

## INTRODUCTION

Shellfish are among the most allergenic classes of seafood and the increase in demand for it has led to more reporting of adverse reactions.^1^ A random telephone survey study reported the prevalence of shellfish or fish allergy was 5.9% of households and was more common in adults (2.8%) compared with children (0.6%).^2^ The prevalence of shrimp allergy in the U.S. was reported to be high as 44-82% of all patients with positive skin prick testing (SPT).^3^ Clinical history, SPT, specific IgE (sIgE) and oral food challenge (OFC) are all used to assess shrimp allergy. Each method has flaws that limit their potential to accurately diagnose shrimp allergy. The history of reaction to shellfish can be confusing as other allergens (seafood and non-seafood) are often concomitantly consumed with shellfish.^1^ Additionally, non-immune mechanisms like exposure of shellfish to marine toxins, viruses, bacteria and parasites, can result in adverse events and confuse documentation of IgE-mediated reactions.^1^ Commercially available shrimp serologic testing does not provide consistent enough specificity or sensitivity to define hypersensitivity or predict clinical reactivity thresholds as have been reported with other food allergens, e.g. peanut or egg. Over 50% of individuals with a history of shrimp allergy will pass a double-blind, placebo-controlled food challenge (DBPCFC), documenting absence of hypersensitivity.^4^ Likewise, the positive predictive value (PPV) for SPT to shrimp is low and limits its use a confirmatory clinical test. NIAID Guidelines for Food Allergy Diagnosis and Treatment promote food challenge with DBPCFC or open OFC as the gold standard for diagnosis.^4–6^ OFC and single blind challenges are excellent tools that may be performed in a clinical setting but are limited by availability in smaller clinical settings and cost.

There is extensive cross-reactivity between different species of seafood. The major allergen in shellfish is tropomyosin and it is responsible for molecular and clinical crossreactivity within the shellfish group and other invertebrates like house dust mite (HDM) and cockroach.^7–9^ Other shrimp allergens such as arginine kinase (AK), myosin light chain (MLC), sarcoplasmic calcium-binding protein (SCP), troponin C and more recently hemocyanin have been identified as potentially clinically relevant allergens.^10–14^ Like tropomyosin, these allergens also are cross-reactive with invertebrate proteins.

The use of component-resolved diagnostics (CRD) with purified natural allergens or recombinant allergens has improved food allergy diagnosis and in some allergen specific settings, like peanut, provided options to predict severity of adverse reactions.^15, 16^ SPT and sIgE to shrimp extract have inferior predictive value compared to sIgE to shrimp tropomyosin.^17^ Even sIgE to shrimp tropomyosin lacks optimal sensitivity (88%) and PPV (71.4%). Although shrimp sensitization has been reported in individuals with concomitant HDM and cockroach sensitization neither invertebrate sensitivity is purported to play a role in allergic reactions noted following shrimp ingestion. In a recent publication, tropomyosin was reported to not likely be a major important allergen in HDM sensitized shrimp allergic patients.^18^ The primary aim of this study was to establish shrimp and component allergen sIgE threshold levels to predict clinical reactivity and diagnose shrimp allergy in individuals with history of adverse reactions to shrimp. The sensitivity and specificity of shrimp component testing in this patient group was also correlated with HDM sensitivity. There have been no published reports to date that categorized shrimp allergic patients based on HDM sensitization and this may be relevant as we try to eliminate HDM cross-reactivity issues.

## MATERIALS AND METHODS

### Patient selection

We have screened 151 patients with diagnosis of shrimp allergy. Of these patients, 79 met inclusion criteria of having a physician diagnosed shrimp allergy with history of positive shrimp SPT and/or elevated shrimp sIgE to shrimp and patients avoiding shrimp because of history of a positive SPT to shrimp and/or elevated sIgE to shrimp. Seventy nine patients were invited and recruited from the Allergy Clinics of the Departments of Pediatric and Internal Medicine in the Sections of Immunology, Allergy and Rheumatology of Baylor College of Medicine, between August 2014 and January 2016. Thirty patients were enrolled, 31 declined OFC and 18 were lost to follow-up (Fig 1). In addition, 10 patients with regular shrimp ingestion and no shrimp sensitization were also recruited as a negative control (NC). The study was approved by Baylor College of Medicine IRB.

### Skin prick test, sIgE antibodies and graded oral challenge

All patients underwent SPT (Greer^®^ Laboratories, Lenoir, NC, USA) to 6 commercial extracts: shrimp, HDM (*Dermatogoides farinae* [*Der f*] and *Dermatogoides pteronyssinus* [*Der p*] and cockroach mix *(Blatella germanica* [*Bla g*] and *Periplaneta americana* [*Per a*]), histamine and negative control. In addition, prick to prick skin test to cooked (boiled for 10 minutes in water) and raw forms of white shrimp (*Litopenaeus vannamei* [*Lit v*]) were also performed. A positive SPT was defined with wheal diameter of at least 3 mm greater than negative control accompanied by erythema 15 minutes after application of extract or fresh forms of shrimp.

Serum IgE antibodies to shrimp (f24, mixture of *Penaeus borealis* [*Pen b*], *Pen m*], *Metapenaeus barbata* [*Met b*] and *Metapenaeopsis joyner* [*Met j*]), HDM (d1, *Der p*) and cockroach (i6, *Bla g*) were measured using ImmunoCAP^®^ (Thermo Fisher Scientific Phadia, Portege, MI). Serum IgE antibodies to native and recombinant allergens were also performed using ImmunoCAP^®^ (*Der p 1* [cysteine protease], *Der p 10* [tropomyosin], recombinant *Penaeus aztecus* [r*Pen a 1*, tropomyosin], native *Penaeus monodon* [n*Pen m 2*, AK], r*Pen m 3* [MLC], r*Pen m 4* [SCP] and r*Pen m 6* [troponin C]). A level ≥ 0.35 kU_A_/L was considered positive.

Of the 30 patients enrolled, 25 patients were consented for an open graded OFC and 5 patients with history of physician-diagnosed anaphylaxis treated with epinephrine, antihistamines, steroids and/or albuterol were enrolled (Fig 1). The shrimp were prepared before the challenge by boiling in water for ≥ 10 minutes. The shrimp was given with doubling increases every 15 minutes starting with 2 g until 32 g dose was reached (total cumulative dose of 62 g). Vital signs (blood pressure, respiratory rate, and pulse rate) and evidence of cutaneous, respiratory, gastrointestinal and cardiovascular signs and symptoms were determined during the challenge. The challenge was discontinued if an objective sign was elicited according to guidelines by the Work Group report on oral food challenge testing.^19^ OFC was performed with emergency medications at bedside in a hospital setting and patients were observed 2 hours after the last dose. Patients were grouped based on the outcome of the OFC.

### Statistical analysis

Demographics, sIgE values and SPT results were compared between using Fisher’s exact test and Wilcoxcon rank sum test. Values >100 kU_A_/L were designated as 100.1 kU_A_/L. Receiver operating characteristic analysis to determine the area under the curve (AUC), Spearman’s correlation, sensitivity and specificity were performed. Differences were considered significant when p ≤ 0.05. The cut-off values for sIgE were determined by choosing the value that reached the highest sensitivity and specificity.

## RESULTS

### Categorization of patients based on OFC and HDM sensitivity

Graded open OFC to shrimp was performed on 25 patients (Table 1). The time reported of patients having a reaction to shrimp before the OFC was between less than one to 45 years and 10 patients have never been exposed to shrimp. Eighteen patients passed the challenge and were classified as shrimp tolerant (ST) patients. Seven patients developed objective signs during the challenge at a median dose of 16 g (range: 2-32 g). Three patients received epinephrine because of severity of symptoms. There were 5 patients in the ST group who developed OAS symptoms that resolved without treatment and completed the OFC without problems. None of the 7 patients who failed the OFC developed biphasic allergic reactions. Five patients did not undergo OFC because of history of physician-diagnosed anaphylaxis upon shrimp ingestion and together with the 7 patients who failed the OFC were classified as shrimp allergic (SA) patients. Patients were further sub-categorized based on their HDM sensitivity, defined as sensitization to major allergen cysteine protease only as this protein is not an allergen in shrimp (*Der p 1* ≥ 0.35 kU_A_/L). SPT to HDM extract was not used as criteria to categorized HDM sensitization as this extract contains HDM tropomyosin and may result in cross-sensitization with shrimp tropomyosin. There were 7 patients in the SA group with HDM sensitization (SA+HDM), 11 patients in the ST group with HDM sensitization (ST+HDM), 5 patients in the SA group without HDM sensitization (SA-HDM) and 7 in the ST group without HDM sensitization (ST-HDM).

**Table 1.**
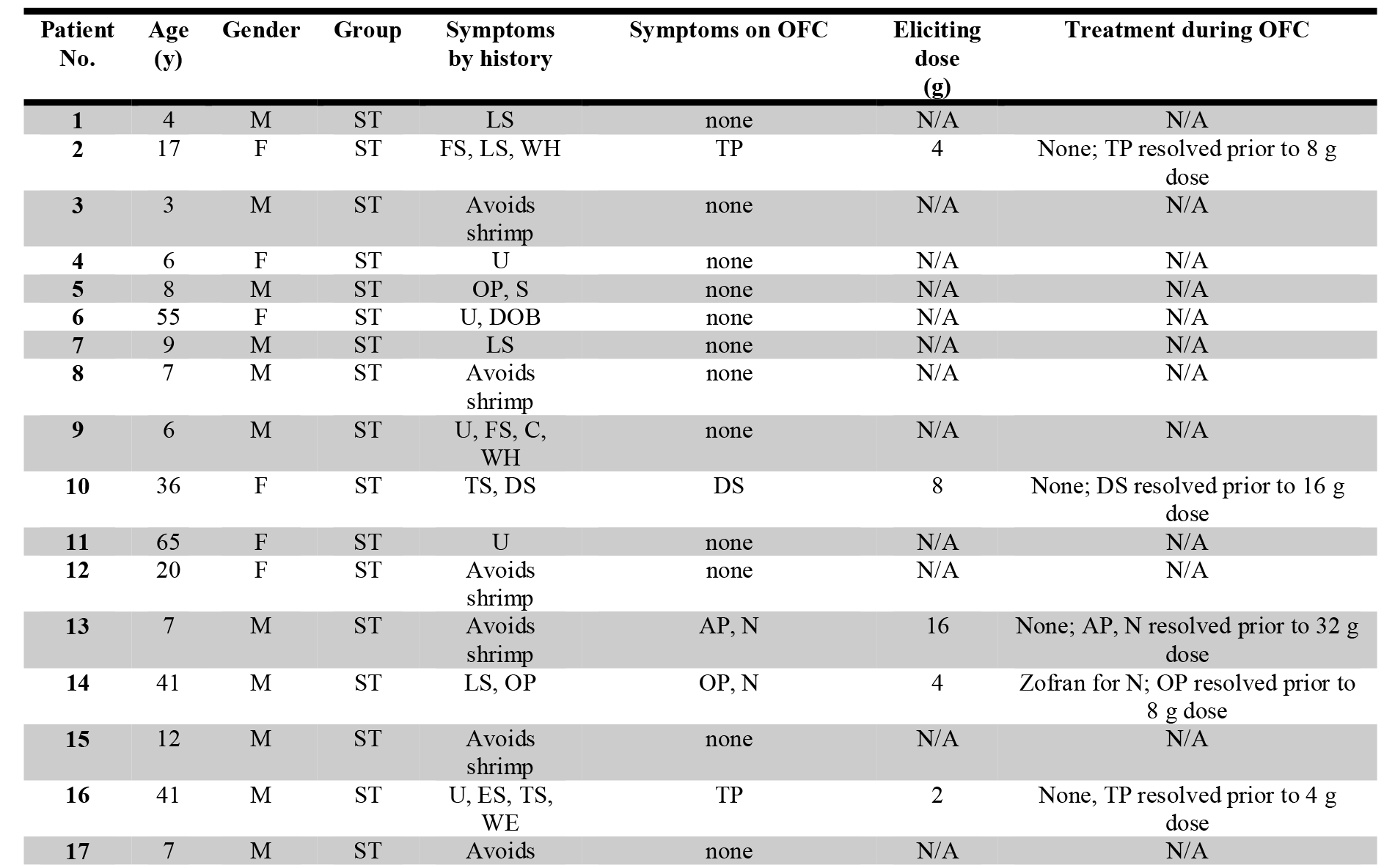
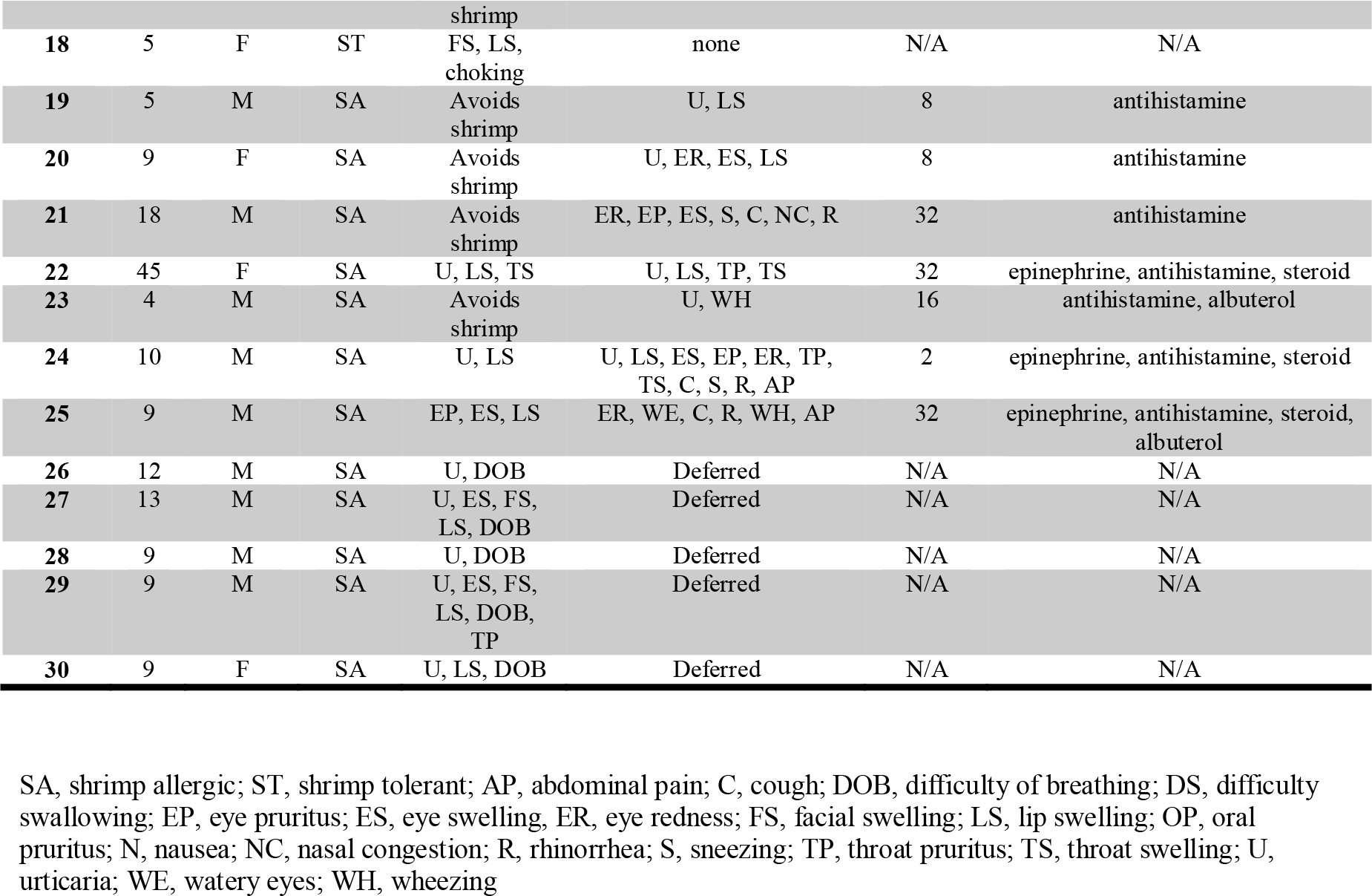
Symptoms by clinical history, outcome of oral food challenge (OFC) and treatment given during OFC.

### Patient characteristics

Patient ages ranged from 4 to 65 years with 20 males and 16 females. The majority of our patients were children (73%). Asthma and other atopic conditions were present in most of patients. The timing of the onset of symptoms was not different between SA and ST (p=0.125). Presence of urticaria was a distinguishing presenting symptom by history in SA compared to SS (p=0.031; Table 2).

**Table 2.** Patient characteristics and symptoms by clinical history.

|  | Shrimp Allergic (SA) | Shrimp Tolerant (ST) | SA vs. ST p-value |
| --- | --- | --- | --- |
|  | 12 | 18 |  |
| Age in years, median (25<sup>th</sup>, 75<sup>th</sup>) | 9 (9, 12.5) | 8.5 (6, 36) | 0.898 |
| Male, N (%) | 9 (75%) | 11 (61%) | 0.694 |
| Race, N (%) |  |  | 0.999 |
| African American | 6 (50%) | 8 (44%) |  |
| Asian | 1 (8%) | 2 (11%) |  |
| Caucasian | 1 (8%) | 2 (11%) |  |
| Hispanic | 4 (33%) | 6 (33%) |  |
| Age at diagnosis in years, median (25<sup>th</sup>, 75<sup>th</sup>) | 6 (3.5, 8.5) | 6 (3, 10) | 0.782 |
| History of asthma, N (%) | 10 (83%) | 13 (72%) | 0.669 |
| History of allergic rhinitis, N (%) | 11 (92%) | 18 (100%) | 0.4 |
| History of eczema, N (%) | 8 (67%) | 10 (56%) | 0.709 |
| History of food allergy other than shellfish, N (%) | 6 (50%) | 9 (50%) | 0.999 |
| History of venom hypersensitivity, N (%) | 0 | 2 (11%) | 0.503 |
| History of allergen immunotherapy, N (%) | 0 | 3 (17%) | 0.255 |
| Onset of symptoms in minutes, N (%) |  |  | 0.125 |
| Immediate (< 10 min.) | 7 (58%) | 4 (22%) |  |
| 10-15 min. | 0 | 4 (22%) |  |
| > 15 min. | 1 (8%) | 4 (22%) |  |
| avoids | 4 (33%) | 6 (33%) |  |
| Urticaria, N (%) | 7 (88%) | 5 (36%) | **0.031** |
| Angioedema, N (%) | 6 (75%) | 7 (50%) | 0.38 |
| Ocular symptoms, N (%) | 2 (25%) | 2 (14%) | 0.602 |
| Respiratory symptoms, N (%) | 4 (50%) | 5 (36%) | 0.662 |
| Oral symptoms, N (%) | 5 (63%) | 6 (40%) | 0.4 |
| Gastrointestinal symptoms, N (%) | 0 | 1 (8%) | 0.999 |

### SPT and sIgE of shrimp allergic and shrimp tolerant patients

SPT positivity was not different between SA and ST groups using shrimp extract (p=0.419), raw shrimp (p=0.605) and cooked shrimp (p=0.221). Of note, 17% of SA group had negative SPT to shrimp extract (2/12) and raw shrimp (1/6), and 38% had negative SPT to cooked shrimp (3/8). However the median wheal diameter to cooked shrimp was larger in SA group than ST group (median wheal, mm [25^th^, 75^th^]: SA=7 [4.5, 13], SS=3 [0, 6], p=0.0294). Similarly, the SPT to *Der p*, *Der f* and cockroach was not different between SA and SS (*Der p*, p=0.999; *Der f*, p=0.26; cockroach, p=0.689; Table 3).

**Table 3.**
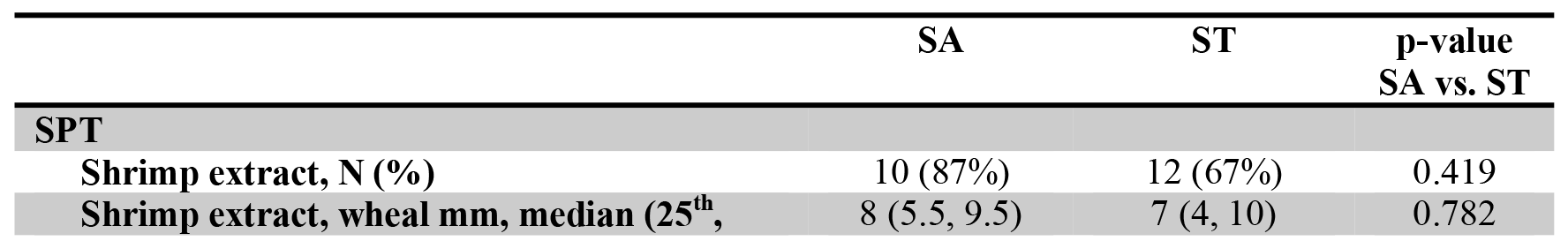
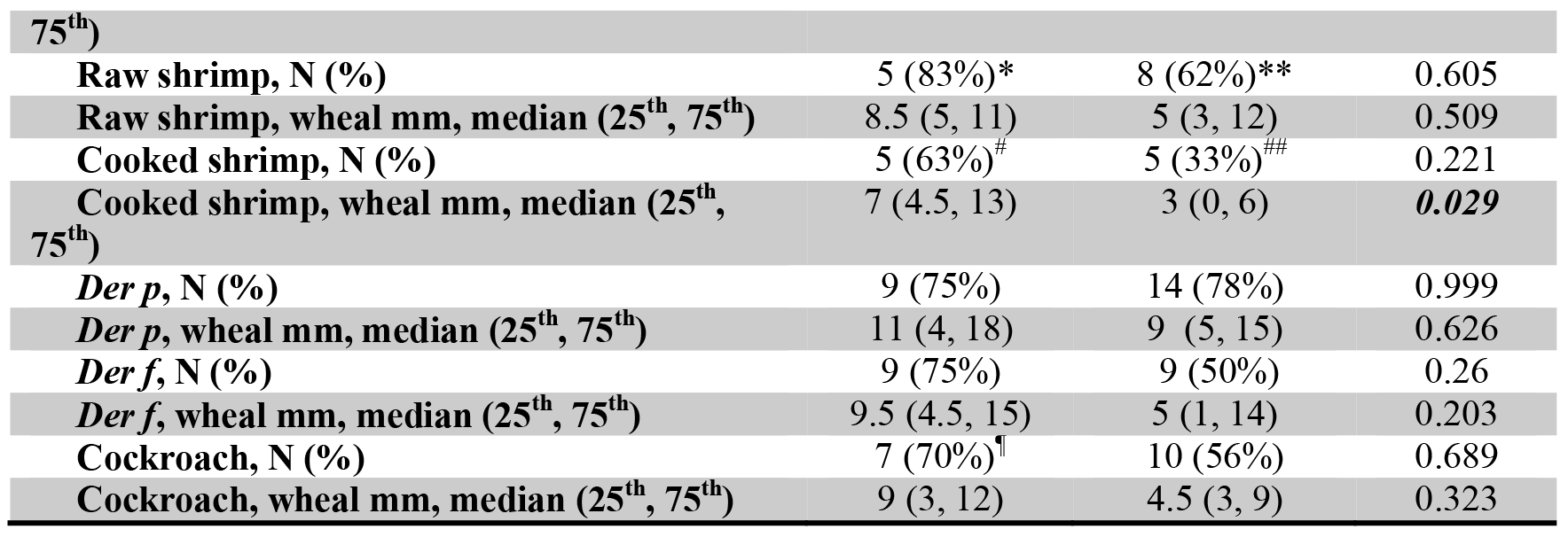
Skin prick test (SPT) results on shrimp allergic (SA) and shrimp tolerant (ST) patients.

The median sIgE to shrimp was elevated in SA (10.2 kU_A_/L) compared to ST (1.25 kU_A_/L) patients (p=0.042). Four patients from ST group (4/18, 22%) and one from SA group (1/12, 8%) had negative sIgE to shrimp and other shrimp proteins. The median sIgE to shrimp and HDM components, and cockroach was not different between groups (Table 4). None of the NC had sensitization to shrimp. There was a trend toward a higher proportion of SA patients with elevated sIgE than ST patients to *rPen a 1* (SA: 83%, ST: 44%, p=0.058) and *Der p 10* (SA: 83%, ST:39%, p=0.026) as shown in Table 4.

**Table 4.** Summary of IgE results of patients enrolled.

|  | SA<br>(n=12) | ST<br>(n=18) | p-value<br>SA<br>vs.<br>ST | SA+HDM<br>(n=7) | ST+HDM<br>(n=11) | p-value<br>SA+HDM<br>vs.<br>ST+HDM | SA-HDM<br>(n=5) | ST-HDM<br>(n=7) | p-value<br>SA-HDM<br>vs.<br>ST-HDM |
| --- | --- | --- | --- | --- | --- | --- | --- | --- | --- |
| sIgE,<br>kU<sub>A</sub>/L |  |  |  |  |  |  |  |  |  |
| Shrimp,<br>N (%)* | 11<br>(92%) | 14<br>(78%) | 0.622 | 6 (86%) | 9 (82%) | 0.228 | 5<br>(100%) | 5<br>(71%) | 0.47 |
| Shrimp,<br>median<br>(25<sup>th</sup>, 75<sup>th</sup>) | 10.2<br>(3.56,<br>13.6) | 1.25<br>(0.47,<br>7.85) | **0.042** | 9.89<br>(1.12,<br>100) | 2.82<br>(0.47,<br>23.6) | 0.365 | 10.5<br>(3.62,<br>11.8) | 1.06<br>(<0.1,<br>3.32) | **0.012** |
| *rPen a*<br>*1*, N (%)* | 10<br>(83%) | 8<br>(44%) | 0.058 | 6 (86%) | 5 (45%) | 0.054 | 4<br>(80%) | 3<br>(43%) | 0.293 |
| *rPen a*<br>*1*, median<br>(25<sup>th</sup>, 75<sup>th</sup>) | 4.8<br>(0.91,<br>8.2) | 0.2<br>(<0.1,<br>4.15) | 0.082 | 4.41 0.72,<br>100) | 0.11<br>(<0.1,<br>15.28) | 0.268 | 5.1<br>(1.1,<br>6.7) | 0.27<br>(<0.1,<br>0.73) | 0.086 |
| *rPen m*<br>*2*, N (%)* | 4<br>(33%) | 7<br>(39%) | 0.999 | 3 (43%) | 6 (55%) | 0.999 | 1<br>(20%) | 1<br>(14%) | 0.999 |
| *rPen m*<br>**2, median**<br>(25<sup>th</sup>, 75<sup>th</sup>) | 0.17<br>($<0.1$,<br>10.8) | 0.14<br>($<0.1$,<br>5.48) | 0.570 | 0.28<br>($<0.1$,<br>33.4) | 0.77<br>($<0.1$,<br>16.8) | 0.544 | 0.11<br>($<0.1$,<br>0.2) | 0.13<br>($<0.1$,<br>0.16) | 0.801 |
| *rPen m*<br>**3, N (%)** * | 3<br>(25%) | 3<br>(17%) | 0.66 | 2 (29%) | 2 (18%) | 0.999 | 1<br>(20%) | 1<br>(14%) | 0.999 |
| *rPen m*<br>**3, median**<br>(25<sup>th</sup>, 75<sup>th</sup>) | $<0.1$<br>($<0.1$,<br>0.84) | 0.06<br>($<0.1$,<br>0.18) | 0.743 | $<0.1$<br>($<0.1$,<br>1.54) | 0.11<br>($<0.1$,<br>0.21) | 0.961 | $<0.1$<br>($<0.1$,<br>$<0.1$) | $<0.1$<br>($<0.1$,<br>0.16) | 0.629 |
| *rPen m*<br>**4, N (%)** * | 4<br>(33%) | 3<br>(17%) | 0.392 | 2 (29%) | 1 (9%) | 0.25 | 2<br>(40%) | 2<br>(29%) | 0.999 |
| *rPen m*<br>**4, median**<br>(25<sup>th</sup>, 75<sup>th</sup>) | $<0.1$<br>($<0.1$,<br>1.02) | $<0.1$<br>($<0.1$,<br>0.12) | 0.405 | $<0.1$<br>($<0.1$,<br>1.17) | $<0.1$<br>($<0.1$,<br>$<0.1$) | 0.619 | 0.13<br>($<0.1$,<br>0.87) | $<0.1$<br>($<0.1$,<br>0.65) | 0.665 |
| *rPen m*<br>**6, N (%)** * | 4<br>(33%) | 1 (6%) | 0.128 | 3 (43%) | 1 (9%) | 0.051 | 1<br>(20%) | 0 | 0.417 |
| *rPen m*<br>**6, median**<br>(25<sup>th</sup>, 75<sup>th</sup>) | $<0.1$<br>($<0.1$,<br>1.32) | $<0.1$<br>($<0.1$,<br>$<0.1$) | 0.209 | $<0.1$<br>($<0.1$,<br>12.9) | $<0.1$<br>($<0.1$,<br>$<0.1$) | 0.188 | $<0.1$<br>($<0.1$,<br>$<0.1$) | N/A | 0.708 |
| *Der p 1*,<br>**N (%)** * | 7<br>(58%) | 11<br>(61%) | 0.999 | 7 (100%) | 11<br>(100%) | 0.622 | 0 | 0 |  |
| *Der p 1*,<br>**median**<br>(25<sup>th</sup>, 75<sup>th</sup>) | 2.8<br>($<0.1$,<br>13.99) | 2.69<br>($<0.1$,<br>48.6) | 0.615 | 9.97 (3.6,<br>28.8) | 38.6<br>(2.81,<br>70.6) | 0.4414 | N/A | N/A | |
| *Der p*<br>**10, N (%)** * | 10<br>(83%) | 7<br>(39%) | **0.026** | 6 (86%) | 5 (45%) | 0.035 | 4<br>(80%) | 2<br>(29%) | 0.242 |
| *Der p*<br>**10, median**<br>(25<sup>th</sup>, 75<sup>th</sup>) | 4.0<br>(1.6,<br>10.99) | 0.2<br>($<0.1$,<br>6.78) | 0.072 | 3.61<br>(0.73, 75) | 0.26<br>($<0.1$,<br>17.4) | 0.363 | 4.09<br>(3.98,<br>6.97) | $<0.1$<br>($<0.1$,<br>0.61) | **0.035** |
| **Cockroach**,<br>**N (%)** * | 11<br>(92%) | 11<br>(65%) | 0.187 | 6 (86%) | 8 (80%) | 0.228 | 5<br>(100%) | 3<br>(43%) | 0.081 |
| **Cockroach**,<br>**median**<br>(25<sup>th</sup>, 75<sup>th</sup>) | 4.92<br>(1.8,<br>8.1) | 1.51<br>(0.19,<br>83.16) | 0.163 | 2.27<br>(0.66,<br>31.7) | 6.48<br>(0.54,<br>12.3) | 0.961 | 6.66<br>(3.66,<br>6.66) | 0.32<br>($<0.1$,<br>1.76) | **0.004** |

### sIgE of shrimp allergic and shrimp tolerant patients based on HDM sensitivity

In patients not sensitized to HDM, median sIgE to shrimp (p=0.012), *Der p 10* (p=0.035) and cockroach (p=0.004) were significantly higher in SA-HDM than ST-HDM patients, while sIgE to *rPen m 2*, *rPen m 3* and *rPen m 4* were not different between the two groups (Table 4). When sIgE was analyzed based on positive HDM sensitivity (*Der p 1 ≥*0. 35 kU_A_/L), the median sIgE levels to shrimp, *rPen a 1* and *Der p 10*trended to be higher in SA+HDM than ST+HDM but this was not significant, while median sIgE to *rPen m 2*, *rPen m3* and cockroach were higher in ST+HDM than SA+HDM patients, again not statistically significant (*rPen m 2* kU_A_/L: p=0.5442; *rPen m 3* kU_A_/L: p=0.961; cockroach kU_A_/L: p=0.961).

### Shrimp sIgE and CRD as a diagnostic test

Since shrimp sIgE was higher in SA than ST patients, the cut-off value that would highly predict clinical reactivity was determined. Currently, commercial sIgE testing have cutoff values set at 0.1 and 0.35 kU_A_/L. Cut-off levels for white shrimp sIgE were determined, having 91.7% sensitivity and 16.7% specificity for 0.1 kU_A_/L, and 91.7% sensitivity and 22.2% specificity for 0.35 kU_A_/L. The AUCs were also not high at both these cut-off levels (0.1 kU_A_/L: 0.54 (0.42, 0.66); 0.35 kU_A_/L: 0.57(0.44, 0.70). When the shrimp sIgE cut-off level of 3.55 kU_A_/L was used in SA patients, it had an AUC of 0.72 (0.53, 0.92), sensitivity of 83.3% and specificity of 66.7%. *Der p 10* sIgE of 2.43 kU_A_/L gave 75% sensitivity and 72.2% specificity, while *rPen a 1* sIgE of 2.48 kU_A_/L had lower sensitivity of 66.7% and specificity of 72.2%. The *Der p 10* AUC was 0.69 (0.49, 0.90) and 0.69 (0.49, 0.89) for *rPen a 1*. Although sIgE cut off values for *rPen a 1* and *Der p 10* had poor sensitivity and specificity, *Der p 10* and *rPen a 1* sIgE levels correlated with each other regardless of HDM sensitivity (r=0.9344).

In non-HDM sensitized SA patients, a shrimp sIgE cut-off level of 3.55 kU_A_/L had AUC of 0.94[0.81, 1.0], sensitivity of 100% and specificity of 85.71%. A *rPen a 1* sIgE of 1.1 kUA/L and *Der p 10* sIgE of 3.98 kU_A_/L both had a high sensitivity, specificity and AUC (Table 5).

**Table 5.** Sensitivity, specificity, receiver operating characteristic (ROC) and area under the curve (AUC) for shrimp, r*Pen a 1* and *Der p 10* IgE levels.

|  | Overall SA vs. ST | SA+HDM vs. ST+HDM | SA-HDM vs. ST-HDM |
| --- | --- | --- | --- |
| **Shrimp IgE cut-off (kUA/L)** | 3.55 | 7.85 | 3.55 |
| **Sensitivity (%)** | 83.3 (51.6, 97.9) | 57.1 (18.4, 90.1) | **100 (47.8, 100)** |
| **Specificity (%)** | 66.7 (41, 86.7) | 54.5 (23.4, 83.3) | 85.7 (42.1, 99.6) |
| **ROC AUC (95% CI)** | 0.72 (0.53, 0.92) | 0.63 (0.34, 0.92) | 0.94 (0.81, 1.0) |
| ***rPen a 1* cut-off (kUA/L)** | 2.48 | 4.41 | 1.1 |
| **Sensitivity (%)** | 66.7 (34.9, 90.1) | 57.1 (18.4, 90.1) | 80 (28.4, 99.5) |
| **Specificity (%)** | 72.2 (46.5, 90.3) | 63.6 (30.8, 89.1) | 85.7 (42.1, 99.6) |
| **ROC AUC (95% CI)** | 0.69 (0.49, 0.89) | 0.66 (0.38, 0.93) | 0.80 (0.47, 1.0) |
| ***Der p 10* cut-off (kUA/L)** | 2.43 | 3.61 | 3.98 |
| **Sensitivity (%)** | 75 (42.8, 94.5) | 57.1 (18.4, 90.1) | 80 (28.4, 99.5) |
| **Specificity (%)** | 72.2 (46.5, 90.3) | 54.5 (23.4, 83.3) | **100 (59, 100)** |
| **ROC AUC (95% CI)** | 0.69 (0.49, 0.90) | 0.63 (0.35, 0.91) | 0.86 (0.57, 1.0) |

## DISCUSSION

Shrimp is one of the less studied food allergens compared to peanut, cow’s milk and egg, and yet it is also one of the most common food implicated in emergency room food-allergic visits^20^. In fact the previously reported rate of anaphylaxis to shrimp is 12% in children and 42% in adults with shrimp allergy.^21, 22^ The natural history of shrimp allergy is less well characterized and it is not known how many affected children will have persistent shrimp allergy into adulthood. Currently, OFC remains the gold standard for the diagnosis of food allergy but OFCs to shrimp is less reported in the literature than OFCs to other foods. This may be due to patient fear due to previous severity of reaction. Hence, less invasive tools are needed to aid clinicians in the diagnosis of shrimp allergy.

The overall rate of positive OFC in this cohort of shrimp allergic patients was 28%, which is lower compared to previous reports (32% to 75%) ^4, 14, 17, 23, 24^ but still higher compared to the overall rate of reactions to OFC in foods in general, which is about 6% in children.^25^ The lower positive shrimp OFC rate in our study may be due to difference in patient population since most of our patients are children. Furthermore, we did not consider symptoms of oral allergy syndrome (OAS) as a failed OFC unless a systemic reaction developed or there were persistent objective signs in the oral mucosa. The prevalence and clinical presentation also varies in different parts of the world. In Asia where there is a higher prevalence of shellfish and HDM allergy, a more common presentation of shrimp allergy is a form of OAS.^24^ There is likely a role of HDM sensitization in development of OAS to shrimp. Of note, the 5 patients in the ST group with OAS that resolved without treatment during OFC had HDM sensitization to cysteine protease and/or tropomyosin, which is likely the primary sensitizer for development of OAS. Although, other HDM allergens are thought to be responsible for HDM and seafood cross-reactivity^26^, it is still unknown if these other allergens may result in OAS. There are no studies to date that have investigated role of HDM sensitization and development of OAS to shrimp.

Shrimp is one of the most common food allergens that clinicians still face a challenge in interpreting SPT and sIgE results as there is still a lack of well-established predictive values that correlate with clinical reactivity. A main limitation is the presence of cross-reacting proteins in shrimp and non-edible arthropods that are common causes of environmental allergies. Shrimp sensitization has been shown to correlate with HDM and cockroach sensitization but may not necessarily relate with clinical reactivity upon shrimp ingestion.^27, 28^ SPT to commercial shrimp extract has low diagnostic efficiency (50.6-65.7%).^17, 24^ Our findings using the commercial shrimp extract and raw shrimp did not distinguish allergic patients from shrimp tolerant patients with shrimp sensitization. However, a wheal size of ≥ 7 mm using cooked shrimp may aid in screening patients suspected of shrimp allergy but must be used with other diagnostic modalities. Cut-off values for SPT wheal size to peanut, egg, sesame and cow’s milk have been established but none for shrimp.^29, 30^ This is the first study that has demonstrated to differential detection of shrimp sensitization in shrimp allergy by SPT wheal size using cooked rather than raw shrimp.

Currently, only shrimp sIgE is available for use clinically and this test alone for diagnosis of shrimp allergy is not reliable because of cross-reactivity among arthropod proteins. Recent reports have shown that sIgE, like SPT, it has low diagnostic efficiency (56.9-74.2%).^17, 24^ Similarly we have shown that although shrimp allergic patients have higher median shrimp sIgE than shrimp tolerant patients, the sensitivity and specificity were only modest. However, a shrimp sIgE cut-off level of 3.55 kU_A_/L has more value in non-HDM sensitized patients as one is not confounded with cross-reactivity issues.

Published reports have been made in improving the diagnosis of shrimp allergy through molecular diagnosis and component resolved diagnostics. IgE epitope recognition to individual shrimp allergens in challenge-proven shrimp allergic individuals have provided modest diagnostic efficiency (*Lit v 1* epitopes 2, 5a and 5b: 80.8%; *Lit v2* epitope 6: 84.6%),^4^ while combination of IgE epitope recognition to tropomyosin and SCP was associated with clinical reactivity (trend to significance, p=0.0549).^14^ Shrimp epitope recognition may also be a useful tool in the future predicting which patients can outgrow shrimp allergy as it was demonstrated that there was greater and more intense epitope recognition of shrimp allergens in children compared to adults.^31^ Although epitope mapping is sensitive, interpretation of results may be difficult since IgE and IgG4 binding areas may largely overlap. Moreover diagnostic efficiency of sIgE to individual shrimp components have also been inconsistent especially for the major allergen tropomyosin (sensitivity 34.2-100%, specificity 45.5-92.8%).^17, 23, 24^ However, it should be noted that the methodology used for determining shrimp tropomyosin differs in the study. In this study, using sIgE to whole shrimp, shrimp (*rPen a 1*) and HDM (*Der p 10*) tropomyosin may potentially be used to diagnose shrimp allergy. Furthermore, the determination of sIgE levels to these proteins are more relevant with higher accuracy in non-HDM sensitized patients since cross-reactivity issues between shrimp and HDM is not a factor anymore. One can also use HDM tropomyosin (*Der p 10*) as a surrogate for shrimp tropomyosin (*rPen a 1*). However, a larger dataset and multi-center study would be needed in the future to confirm findings and potentially applying it in clinical practice. Majority of shrimp allergic patients regardless of HDM sensitivity have a greater proportion of their sIgE bind to shrimp tropomyosin (80-86%) compared to the other shrimp proteins, which suggests that it is the major relevant shrimp allergen. IgE to other shrimp proteins like AK (n*Pen m 2*), MLC (*rPen m 3*), SCP (*rPen m 4*) and troponin C (*rPen m 6*) were not more frequent nor higher in shrimp allergic patients compared to tropomyosin possibly due to small number of patients.

## CONCLUSIONS

The use of shrimp and CRD sIgE is helpful in predicting clinical reactivity in shrimp allergic patients who are not HDM sensitized. A shrimp sIgE level that is 100% sensitive (≥3.55 kUA/L) and a *Der p 10* sIgE level that is 100% specific (≥3.98 kUA/L) may be utilized to confirm shrimp allergy in non-HDM sensitized shrimp allergic patients. *rPen a 1* may also be used in these subset of patients with moderate sensitivity (80%) and specificity (85.7%). SPT to shrimp extract is not a reliable tool in screening for shrimp allergy. The use of wheal size diameter to cooked shrimp SPT is more useful than the extract to diagnose clinical shrimp allergy. In summary, we provided evidence on the limitation of the component resolved diagnostics in shrimp as it shares homologous proteins with dust mite and cockroach. However, it may be of value in patients with no dust mite or cockroach sensitization as cross-reactivity is not a factor anymore when interpreting the results.

## Acknowledgements

This study was supported by Thermo Fisher Scientific Phadia.

